# Single molecule detection of Tau seeds in extracellular vesicles from Alzheimer’s disease brains

**DOI:** 10.64898/2026.09.21.753085

**Authors:** Dalya Gulseren, Anaïs Bécot, The brain bank NeuroCEB Neuropathology Network, Christian G Specht, Mehdi Kabani

## Abstract

Extracellular vesicles (EVs) play a critical role in the propagation of Tau pathology in Alzheimer’s disease (AD). The selective identification and characterization of Tau seed-carrying EVs would help developing new diagnostic tools and therapeutic strategies. While pathogenic forms of Tau can be detected biochemically in EV-enriched fractions, only super-resolution fluorescence imaging provides both morphological and compositional information at the level of single EVs on account of their small size and the low number of target molecules. This was the object of our study. Post-mortem human brain-derived EVs from AD patients and non-demented control individuals were characterized in depth using biochemistry, transmission electron microscopy (TEM) and single molecule localization microscopy (SMLM). Dual-colour SMLM of immobilized EVs, displaying size distributions that were consistent with TEM, enabled statistically meaningful measurements of thousands of EVs in parallel. Owing to the high spatial resolution and capability of SMLM to detect sparse epitopes, we identified pathological Tau proteins in individual EVs from AD samples. Reconstructed super-resolution images of permeabilized EVs demonstrate that pathological Tau forms are frequent cargo proteins of EVs from AD brains. Our data illustrate the power of SMLM to determine the ultrastructure, molecular composition and topology of single EVs and EV populations.

---

Alzheimer’s disease (AD) is the most prevalent form of dementia. It is characterized by the misfolding and aggregation of amyloid-β (Aβ) peptides and hyperphosphorylated Tau proteins into amyloid fibrils(Fitzpatrick et al. 2017; Yang et al. 2022). While the former accumulates as extracellular plaques, hyperphosphorylated Tau forms intraneuronal inclusions known as neurofibrillary tangles and neurites(Braak et Braak 1991). The spreading of Tau pathology follows stereotypical patterns referred to as Braak stages and correlates well with neurodegeneration and cognitive decline in AD patients(Braak et Braak 1991; Arriagada et al. 1992). This process is thought to be mediated by a prion-like propagation mechanism where pathological Tau seeds are transmitted to unaffected cells, where they act as templates for the aggregation of soluble Tau protein(Colin et al. 2020). Several mechanisms were proposed to account for the cell-to-cell transmission of Tau seeds, including the release and uptake of free seeds by exocytosis and endocytosis(Frost et al. 2009; Holmes et al. 2013), respectively, as well as passage via nanotubes(Chastagner et al. 2020). In recent years, extracellular vesicles (EVs) emerged as major drivers of Tau propagation in AD and other tauopathies(Leroux et al. 2022; Ruan et al. 2021; Wang et al. 2017).

EVs are lipid bilayer-enclosed nanosized particles released by most cells, including neurons and glia in the central nervous system(Chen et al. 2025). EVs are heterogeneous in size, molecular composition and functions and are broadly categorized into either exosomes or ectosomes according to biogenesis(van Niel et al. 2022). Exosomes, ranging from 30 to 150 nm in diameter, originate from endosomal compartments and are secreted via exocytosis following the fusion of multivesicular bodies (MVB) with the plasma membrane. Ectosomes, ranging from 50 nm to several microns in diameter, are formed via the outward budding and fission of the plasma membrane(van Niel et al. 2022). EVs are but a subset of all extracellular particles that can be secreted or shed by cells, such as apoptotic bodies, oncosomes, exophers or migrasomes(Buzas 2023; Welsh et al. 2024), highlighting the importance of proper isolation and characterization methods(Welsh et al. 2024). EVs carry a wide range of luminal or membrane-associated proteins, nucleic acids, lipids and metabolites, which they can deliver to the cytosol of recipient cells (van Niel et al. 2022). EVs are not merely a cellular waste disposal system as thought initially(Johnstone et al. 1987), but have been shown to play a role in cell-cell communication and the regulation of biological processes, both in physiological and pathological contexts(van Niel et al. 2022). In the central nervous system, EVs are thought to help maintain brain homeostasis and promote neuroprotection, but also to act as vehicles for the intercellular transmission of pathological proteins (*e.g.* Tau, α-synuclein) in neurodegenerative diseases, (*e.g.* Alzheimer’s disease, Parkinson’s disease)(Chen et al. 2025; Colin et al. 2020). Furthermore, brain EVs have the potential to cross biological barriers, including the blood-brain barrier(Krämer-Albers 2022). They can be found in biofluids, such as plasma and cerebrospinal fluid (CSF), which makes them a promising source of biomarkers.

Hyperphosphorylated Tau assemblies are found in EV fractions isolated from brain tissue or CSF of AD patients(Ruan et al. 2021; Saman et al. 2012; Crotti et al. 2019; Miyoshi et al. 2021; Leroux et al. 2022; Fowler et al. 2025; Tyagi et al. 2026). These EV-associated Tau seeds can induce Tau pathology in cellular models and animals(Ruan et al. 2021; Crotti et al. 2019; Miyoshi et al. 2021; Leroux et al. 2022). Seeding-competent Tau species were described both in small and large EVs(Ruan et al. 2021; Leroux et al. 2022; Fowler et al. 2025; Dujardin et al. 2014; Oosterlynck et al. 2025; Tyagi et al. 2026). However, the association of Tau with EVs has been mostly done using biochemical approaches in bulk EV preparations or using electron microscopy (EM) coupled with immunogold labeling(Crotti et al. 2019; Leroux et al. 2022; Ruan et al. 2021; Saman et al. 2012; Tyagi et al. 2026). Neither of these methods provide ultrastructural and topological information about which Tau species are associated with EVs and how. In a recent study, bundles of Tau filaments were visualized for the first time within AD brain-derived EVs using cryo-electron tomography(Fowler et al. 2025). However, these observations were restricted to very large (>400-600 nm in diameter) and low abundant EVs that often contained multiple smaller vesicles within their lumen(Fowler et al. 2025). These characteristics are reminiscent of multivesicular bodies or apoptotic bodies resulting from neuronal cell death rather than those of EVs. In support of these observations however, human induced pluripotent stem cell (iPSC)-derived microglia-like-cells fed with recombinant Tau fibrils were shown to secrete EVs packed with these Tau fibrils, as demonstrated by cryo-electron tomography(Karabova et al. 2026). Importantly, the diameter of the EVs correlated with the length of the fibrils they contained(Karabova et al. 2026). A separate study found no Tau filaments but only Tau-positive globular structures, which were hypothesized to be Tau oligomers or protofibrils, in sarkosyl-insoluble fractions of AD brain-derived EVs(Ruan et al. 2021). Hence, the identity and molecular properties of the EVs responsible for Tau propagation are still unclear.

Deciphering specific signatures of EVs containing Tau seeds could not only improve our knowledge of the pathophysiology of AD, but could also lead to new diagnostic and prognostic tools(Chen et al. 2025). This requires sensitive and quantitative methods that give access to precise ultrastructural, topological, and compositional information about individual EVs as well as EV subpopulations(Alexandre et al. 2025). Among different techniques (*e.g.* EM, flow cytometry) allowing multiparametric single-vesicle analysis, single molecule localization microscopy (SMLM) is particularly well-suited for EV profiling, due to its exceptional sensitivity and spatial resolution at the nanometer scale(Welsh et al. 2024; Alexandre et al. 2025). In the present study, we identify pathological Tau in intact EVs isolated from post-mortem AD cortical tissue samples using dual-colour SMLM, demonstrating the power of single molecule super-resolution microscopy as a sensitive analytical tool for biomedical research.

## Results

### Isolation of EVs from post-mortem AD and control brain tissues

We isolated EVs from post-mortem frontal and parietal cortex tissue from patients with AD (Braak stages V/VI) as well as from age-matched non-demented individuals (Supplementary Table S1). The ability to isolate intact EVs with preserved protein topology and organization is critical in order to assess their molecular composition and ultrastructural properties. To achieve this, the brain tissue pieces were gently dissociated with limited mechanical processing and using collagenase D (see Materials and Methods), as this enzyme was shown to preserve the integrity of EVs with only minimal proteolytic cleavage of vesicle surface-exposed proteins compared to other enzymes such as papain(Oosterlynck et al. 2025; Matamoros-Angles et al. 2024). After removing cells and tissue debris by low-speed centrifugation and filtration, EVs were isolated from the resulting interstitial fluid using size exclusion chromatography (SEC). To preserve at best the integrity of the EVs, and because Tau seeds may be found in both small and large EVs(Dujardin et al. 2014; Fowler et al. 2025; Leroux et al. 2022; Oosterlynck et al. 2025; Ruan et al. 2021), we did not further separate EVs based on their size or density.

Brain-derived EV preparations were visualized by negative-staining transmission electron microscopy (TEM), which revealed the presence of vesicles with the expected ovoid, round or cup-shaped morphologies, the latter likely resulting from dehydration during sample preparation (Fig. 1A, Supplementary Fig. S1). The size distribution of EVs was strongly skewed towards smaller vesicles, with a median size of 55±2 nm and 60±2 nm for AD and control samples, respectively. Their diameters ranged from 35 to 250 nm, with a small fraction (less than 2%) of larger vesicles of up to 600 nm (Fig. 1B). We then characterized EVs using tunable resistive pulse sensing (TRPS) analysis(de Vrij et al. 2013). Based on these TRPS measurements, the concentrations of AD and control EVs, normalized to the weight of processed brain tissue, were within similar range, at 10^11^ ± 10^10^ particles/mL/g (Fig. 1C). The size distribution of 91 ± 2% of AD and control EVs measured by TRPS ranged from 85 to 250 nm in diameter (Fig. 1D), as the technique imposes a lower cut off. Larger vesicles of up to 500 nm accounted for 9 ± 2% of the total (Fig. 1D). In agreement with previous studies(Leroux et al. 2022; Ruan et al. 2021), we did not observe significant differences in the size (Fig. 1B,D; p≈1, Kolmogorov-Smirnov test) or concentration (Fig. 1C; p=0.4857, two-tailed Mann-Whitney) of AD and control EVs.

**Figure 1.**
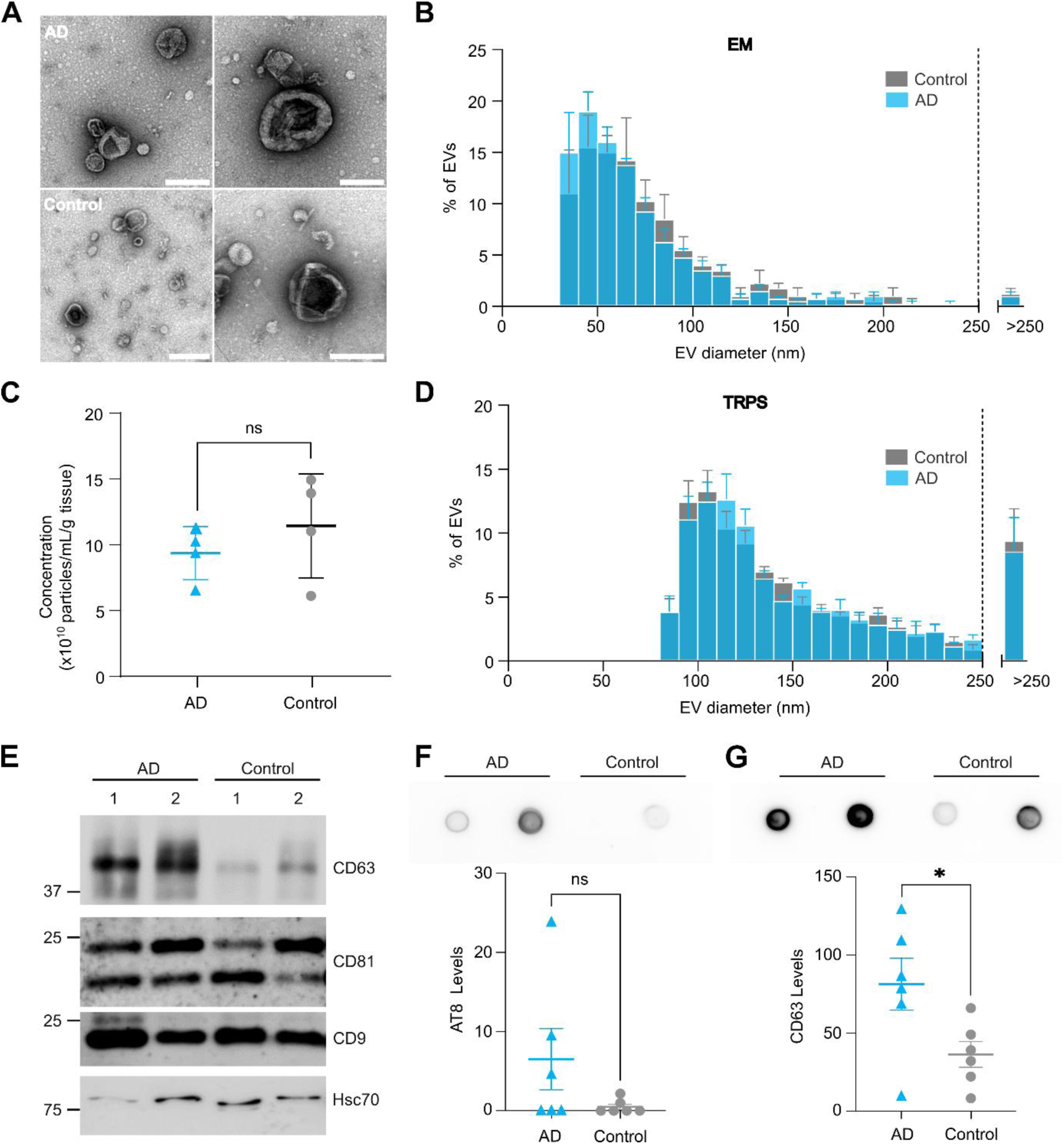
Isolation and characterization of brain-derived EVs. **(A)** EVs isolated by size exclusion chromatography (SEC) from AD (upper panels) or control (lower panels) tissues were visualized by negative-staining transmission electron microscopy (TEM). Representative panels of EVs of varying sizes (each from the same individual) are shown (scale bar: 200nm). Additional images can be found in Supplementary Fig. S1. **(B)** Representative particle size distribution histograms of EVs from AD (blue bars) or control (gray bars) brains determined by TEM (only particles larger than 35 nm were taken into account; N=4 AD; N=4 control; 200 particles measured/sample; data represent mean ± SEM). **(C)** Particle concentration of EVs from AD (blue dots) or control (gray dots) tissues determined by tunable resistive pulse sensing (TRPS) using an NP200 nanopore (N=4 AD; N=4 control; 2 replicates/sample; data represent mean ± SEM; two-tailed Mann-Whitney test). **(D)** Representative particle size distribution histograms of EVs from AD (blue bars) or control (gray bars) brains determined from the TRPS data in (C). Data were normalized to the total number of particles set at 100%, and EVs larger than 250 nm were represented as a single subpopulation (after the dotted line; >250). (N=4 AD; N=4 control; 2 replicates/sample; data represent mean ± SEM). **(E)** Western Blot analysis of surface (CD9, CD63, CD81) and intraluminal (Hsc70) EV markers in EVs from AD (left lanes) or control (right lanes) tissues. Equal volumes of EVs were loaded for all conditions. **(F)** Quantification of AT8-positive Tau levels in EVs from AD or control brains by dot blot analysis (equal amounts of EVs were spotted; N=6 AD; N=6 control; two-tailed Mann Whitney). A representative dot blot is shown (top panel). **(G)** Quantification of CD63 levels in EVs from AD or control brains by dot blot analysis (equal amounts of EVs were spotted; N=6 AD; N=6 control; two-tailed Mann Whitney). A representative dot blot is shown (top panel).

We then confirmed by Western blot the presence of the surface-exposed tetraspanins CD9, CD63 and CD81, as well as intraluminal Hsc70, all of which are expected EV markers (Welsh et al. 2024) (Fig. 1E). We observed significantly higher CD63 levels in EVs from AD patients compared to control ones (Fig. 1E,G). In previous studies reporting the characterization of AD brain-derived EVs, CD63 was either not detected(Leroux et al. 2022; Oosterlynck et al. 2025) or its levels not assessed quantitatively(Ruan et al. 2021). However, elevation of CD63 levels in brain-derived EVs has been reported previously in association with aging or under other disease conditions in mice and humans, and has been attributed to increased release of exosomes(Gauthier et al. 2017; Kim et al. 2022).

To detect pathological Tau in our EVs, we used the AT8 antibody which is widely used to detect the paired-helical filaments (PHF) of Tau, which are the main components of neurofibrillary tangles found in AD(Goedert et al. 1995). Tau phosphorylation at residues S202 and T205 is required for recognition by AT8, with binding enhanced by phosphorylation at S208(Malia et al. 2016). AT8-positive Tau was detected in varying amounts in all EV samples, with those from AD patients generally displaying higher levels than control ones (Fig. 1F). It is important to note that AT8 can also detect soluble pre-fibrillar Tau species at early stages of AD, as well as phosphorylated Tau proteins in the normal adult brain(Goedert et al. 1993; Koss et al. 2016; Matsuo et al. 1994). The presence of AT8-positive Tau in EVs isolated from control individuals is therefore expected and could be due to aging, undiagnosed pathologies or the aforementioned reasons. Immunogold labeling EM further confirmed that AT8-positive Tau is associated to EVs in AD samples (Supplementary Fig. S2).

### Dual colour super-resolution microscopy of brain-derived EVs

To further characterize our brain-derived EVs at single-vesicle level, we applied super-resolution imaging based on single molecule localization microscopy (SMLM). To this aim, we set up simultaneous two-colour SMLM using spectral demixing of two far-red fluorophores (AF647, CF680) with overlapping emission spectra (Fig. 2). The strength of this approach lies in its sensitivity and the capacity to detect specific molecular cargo within individual EVs. The SMLM experiments were performed by capturing EVs without fixation or permeabilization on coverslips functionalized with lectins, which bind to EV surface-exposed glycans (see Materials and Methods). Captured EVs were then labeled with a cocktail of antibodies that target tetraspanins CD9, CD63 and CD81 and that were conjugated with Alexa Fluor 647 (AF647) fluorophore. Alternatively, individual tetraspanins were labeled using specific primary antibodies and CF680-conjugated secondary antibodies.

**Figure 2.**
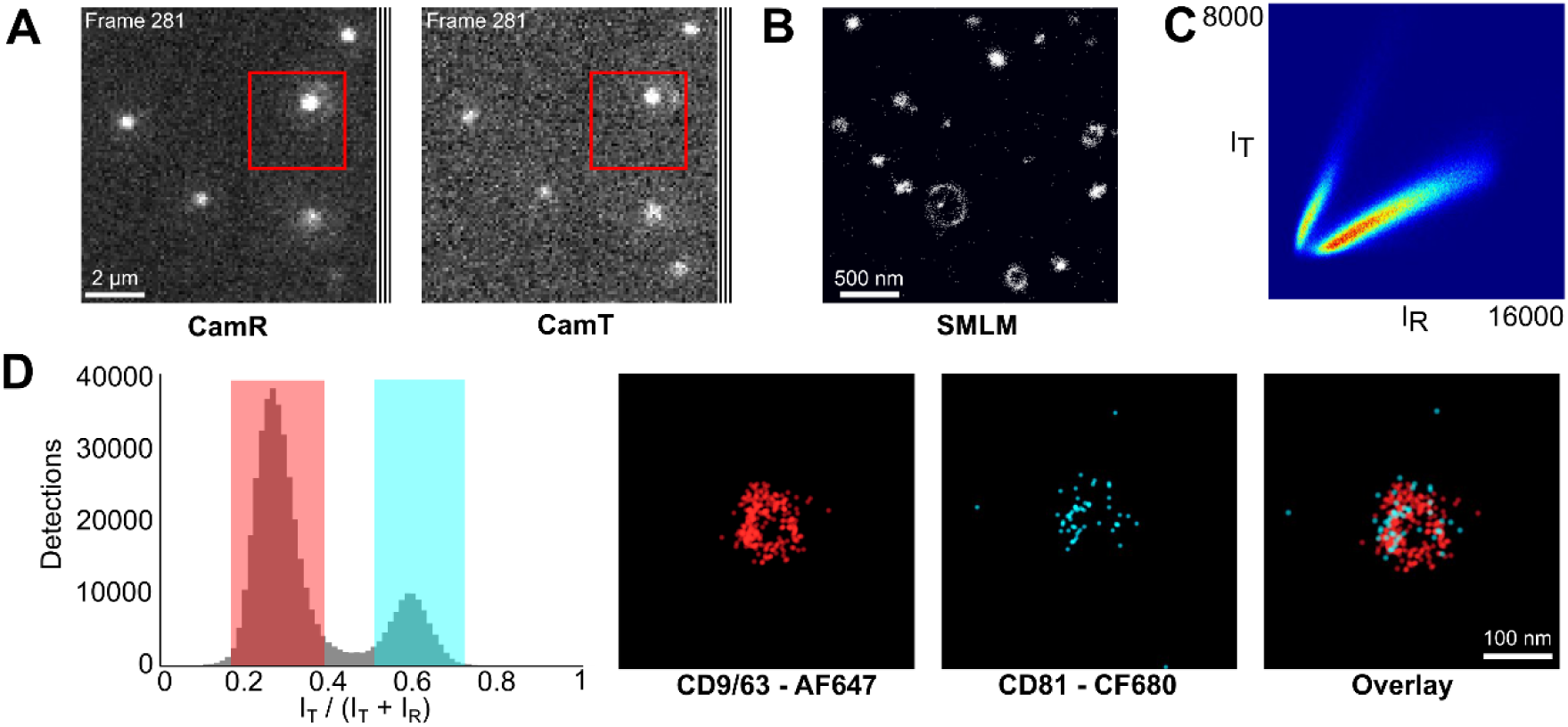
Dual colour super-resolution microscopy of brain-derived EVs. **(A-D)** Spectral demixing was performed on brain-derived EVs labelled with CD9 and CD63 coupled to AF647, and CD81 coupled to CF680. SMLM recordings of 20000 frames were taken in spectral demixing mode. **(A)** Raw image frame (No. 281) acquired simultaneously with the reflected (CamR) and the transmitted camera (CamT). Scale bar: 2 µm. **(B)** Pointillist SMLM image of a zoomed region (red square in A), showing the combined detections of both channels (before demixing). Scale: 500 nm. **(C)** Photon map of single molecule detections shows two distinct populations of intensities (in arbitrary units) corresponding to the AF647 (IR) and CF680 fluorophores (IT). **(D)** On the left, the intensity ratio histogram of single molecule detections is shown in gray, as well as the intensity ranges chosen to separate the AF647 detections (red, 0.18-42) and the CF680 detections (cyan, 0.48-0.74). Right: Super-resolution pointillist images of a single EV displaying both CD9/63 detections (in red), and CD81 detections (cyan). The overlay shows the co-localization of both markers in the same EV.

SMLM recordings of 20000 frames were acquired at 20 Hz using a 640 nm laser, adding low intensity UV illumination to maintain efficient blinking of the fluorophores (see Materials and Methods). For spectral demixing, a 700 nm beam splitter was placed in the emission path to preferentially direct the AF647 and the CF680 signals towards the reflected (CamR) and the transmitted (CamT) camera, respectively (Fig. 2A). Given the overlapping emission spectra, detections from both fluorophores are represented in cumulative pointillist images (Fig. 2B) that can be separated by calculating their intensity ratios in the two channels (Fig. 2C).

Rendered SMLM images of EVs labelled using different combinations of tetraspanin antibodies show round clusters of detections of different sizes, typically in the order of 100 nm (Fig. 2D and Supplementary Fig. S3). Large vesicles often appear as halos due to the projection of the single molecule detections in two-dimensional recordings (Fig. 2D and Supplementary Fig. S3A), whereas smaller ones are seen as a dense cloud of detections (Supplementary Fig. S3B). Dual-colour SMLM further shows the colocalization of CD9 and CD63 (labelled with AF647) and of CD81 (labelled with CF680) within the same vesicles (Fig. 2D). The specificity of fluorophore demixing was confirmed through control experiments in which all three tetraspanins were labelled either with AF647 or with CF680. In these recordings, only a single peak of intensity ratio is apparent, with only few incorrectly attributed detections (Supplementary Fig. S3). These results demonstrate the feasibility of two-colour super-resolution imaging of highly proximal EV proteins with minimal interference.

### Detection of pathological Tau in EVs from AD brain

Our data show that brain-derived EV preparations of AD patients contain phosphorylated Tau protein (Fig. 1). Given that SMLM imaging has the resolution and sensitivity to visualize EVs at the single vesicle level, we analyzed EVs prepared from brain tissue of AD patients and of non-demented control individuals to characterize their hyperphosphorylated Tau cargo.

EVs from AD (N=5 patients) and control brains (N=4 individuals) that had not been fixed or permeabilized were captured on lectin-functionalized coverslips, and labeled as before using AF647-conjugated antibodies against CD9, CD63, and CD81. AT8 antibodies and CF680-conjugated secondary antibodies were used to detect Tau. Dual colour acquisition was then performed using spectral demixing, and the intensity ratios for each fluorophore determined (Fig. 3A,B). A large peak of AF647 detections was observed for both AD and control EVs, which after cluster analysis confirmed the presence of ∼2500 EVs per field of view (FOV, 2496 ± 672, n=27). The overall size distribution of the EVs was similar to that determined by EM analysis of the initial samples (Fig. 1A,B). However, CF680 detections were very scarce in both AD and control EVs (Fig. 3A,B) which could be due to low levels of AT8 epitopes in these samples, or their inaccessibility to the antibodies (see below).

**Figure 3.**
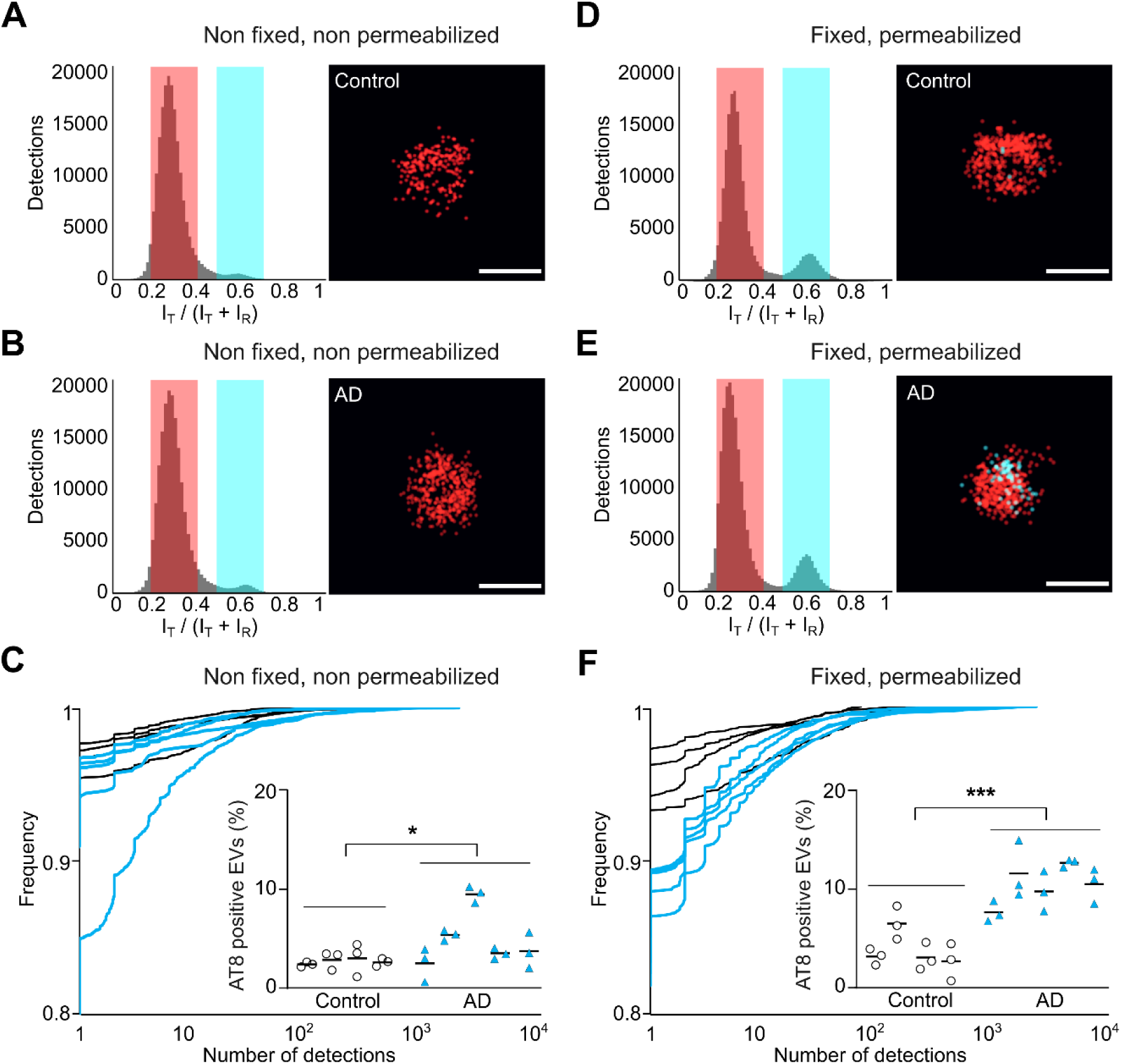
Detection of AT8-positive Tau in control and AD brain-derived EVs by SMLM. **(A, B)** Dual colour SMLM using spectral demixing was carried out with EVs (no fixation or permeabilization) labelled with CD9/63/81 coupled to AF647. Left: Intensity ratio histograms of detections corresponding to CD9/36/81 (red) and phoshorylated Tau (AT8, cyan) in EV fractions from **(A)** control and **(B)** AD brains. Right: representative pointillist images of EVs (scale bars: 100 nm). **(C)** Quantification of the number of AT8 detections per EV represented as cumulative distribution in the non-permeabilized control (black traces) and AD patient samples (cyan traces). The insert shows the fraction of AT8 positive EVs in three separate recordings and the mean value for each EV preparation (n = 12 recordings from 4 controls and n = 15 recordings from 5 AD patients; * p = 0.0034, Mann-Whitney test). **(D, E)** Dual colour SMLM was carried out with the same EV samples after fixation and permeabilization. Left: intensity ratio histograms of **(D)** control and **(E)** AD brains labelled with anti-CD9/63/81 (AF647, red) and AT8 (AT8, CF680, cyan) antibodies. Right: representative pointillist images of single EVs (scale bars: 100 nm). **(F)** Quantification of the AT8 detection number per EV shown as cumulative distribution of the EV population in permeabilized samples (black traces for controls, cyan for AD patients). Insert: Fraction of AT8-positive EVs (≥ 2 detections) for each recording and each sample (nctrl = 12, nAD = 15, *** p < 0.0001, Mann-Whitney test).

To quantify the AT8 signals associated with the EVs, we displayed the number of AT8 detections per EV as a cumulative distribution for AD and control EV samples (Fig. 3C). Even though the majority (>95%) of EVs recorded under these conditions are devoid of AT8 detections, we observed a slight rightward shift of the distribution of AT8 detections in most AD samples compared to controls. We also calculated the fraction of EVs containing at least 2 AT8 detections in 3 separate recordings for each sample (Fig. 3C, insert). The proportion of AT8-positive AD EVs was significantly elevated in the AD samples compared to the control group, with a p-value of 0.0034 (n_ctrl_ = 12 recordings from 4 control brains, n_AD_ = 15 recordings from 5 AD patient brains, two-tailed Mann-Whitney test). In absolute terms, however, the distribution of the number of AT8 detections and the fraction of positive EVs was only marginally increased in the AD samples (about two-fold compared to the control).

In the experiments described above, the EV membrane is assumed to be essentially intact. Because Tau is thought to be an intraluminal EV cargo(Dujardin et al. 2014; Fowler et al. 2025; Leroux et al. 2022; Ruan et al. 2021), AT8 antibodies likely detect Tau molecules present at the EV surface. In order to improve access of the antibodies to the lumen of the EVs, we adapted earlier permeabilization protocols(Kabani et al. 2020; Kabani et Melki 2015). EVs were captured on lectin-functionalized coverslips as before, but were then fixed with paraformaldehyde and gently permeabilized with Triton-X100 at low concentration before labeling (see Materials and Methods). EM analysis confirmed that fixation and permeabilization did not alter EV morphology (Supplementary Fig. S4), even though the total number of vesicles in SMLM was reduced (1374 ± 282 EVs per FOV, n=27) and the size distribution slightly altered (Fig. 4B). Under these conditions, a greater number of CF680 detections were recorded, giving rise to a higher CF680 peak in the intensity ratio histograms (Fig. 3D, E). This was likely due to the improved access of AT8 epitopes after permeabilization, both in the AD and control samples.

**Figure 4.**
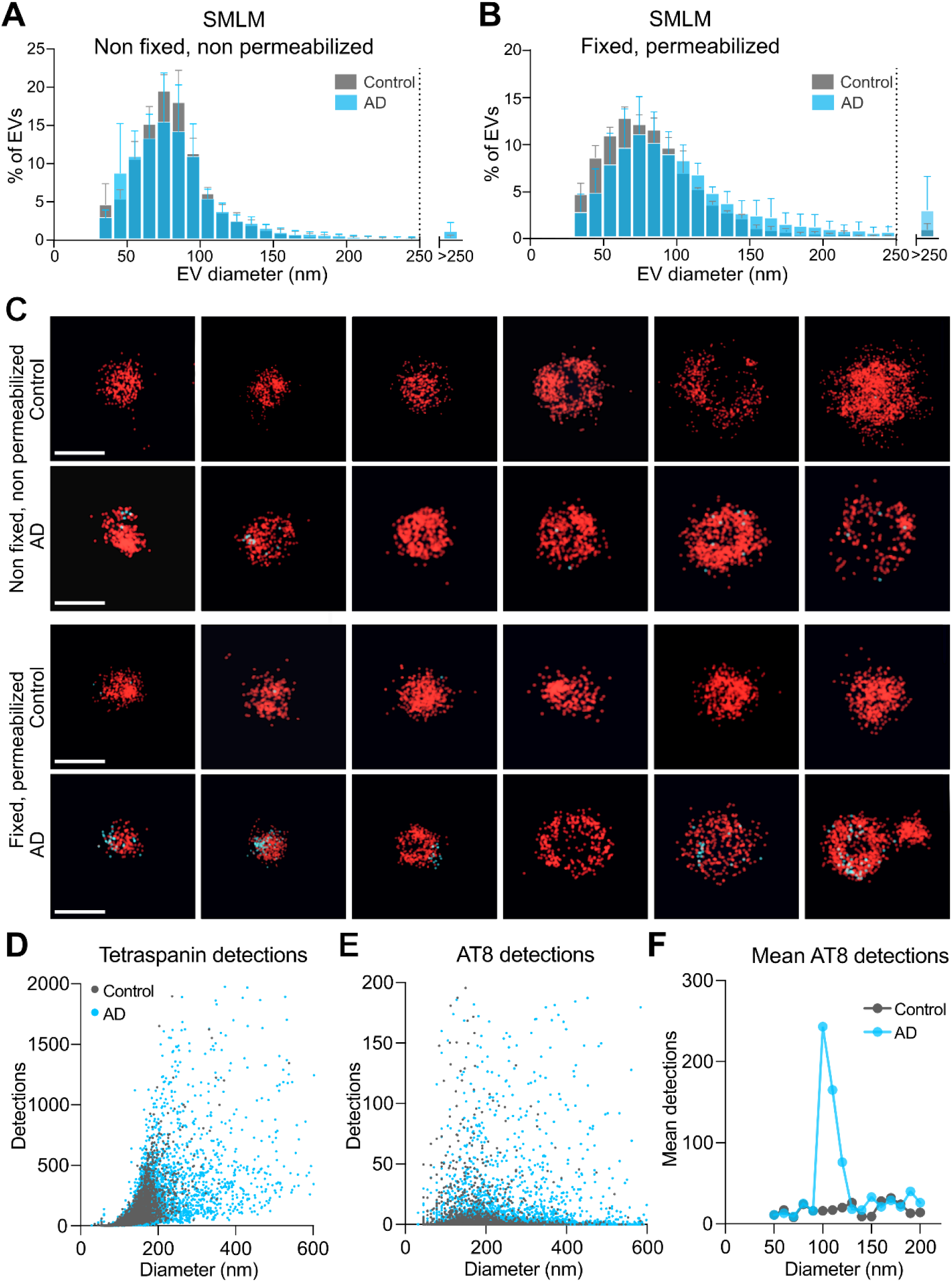
Quantitative and qualitative SMLM-based characterization of Tau-containing EVs. Dual-colour SMLM was carried out with EVs labelled with anti-CD9/63/81 (AF647, red) and anti-Tau antibodies (AT8, CF680, cyan), with or without fixation and permeabilization. **(A, B)** Particle size histograms of EVs from AD (cyan bars) or control (gray bars) brains as determined by SMLM (same dataset as in Fig. 3; mean ± SEM). Data were normalized to the total number of particles set at 100%, and EVs larger than 250 nm were represented as a single bin (beyond the dotted line; >250). **(C)** Representative pointillist images of single EVs of increasing sizes (see also Supplementary Fig. S5). **(D,E)** Number of tetraspanin detections (CD9, CD63, CD81) and AT8 detections plotted as a function of EV diameter. **(F)** The mean number of AT8 detections per EV was calculated for the EVs with the strongest AT8 signals (top 5% of the population in both datasets) and shown according to size.

At the level of individual EVs, we observed that all samples had vesicles associated with clusters of AT8 detections, but this was more evident in the super-resolution image reconstructions of AD samples (Fig. 3E, see also Fig. 4C, Supplementary Fig. S5). When the number of AT8 detections per EV was represented as cumulative distribution, we observed a marked rightward shift in the AD group compared to the control group (Fig. 3F). The percentage of AT8-positive EVs was also significantly higher in the AD samples than in the controls (Fig. 3F, inset), with a p-value of <0.0001 (n_ctrl_ = 12 FOVs from 4 control brains, n_AD_ = 15 FOVs from 5 AD brains, two-tailed Mann-Whitney test).

At the population level, we did not observe any significantly differences in the size distribution of EVs from AD and control brains. This was the case for both the native EVs (Fig. 4A; p≈1, Kolmogorov-Smirnov test), as well as the fixed and permeabilized samples (Fig. 4B; p≈1, Kolmogorov-Smirnov test). In order to assess whether AT8-positive Tau was specifically enriched in particular fractions of EVs, we plotted the number of CD9, CD63 and CD81 detections and AT8 detections as a function of EV diameter (n=15 FOVs from 5 AD patients and n=12 FOVs from 4 control individuals, Fig. 4D-E). As expected, we observed a positive correlation between the tetraspanins detections and EV size (Fig. 4D). This was not the case for AT8 detections that displayed a scattered distribution with no obvious correlation with EV size (Fig. 4E). Nevertheless, the analysis confirmed that the number of AT8 detections was higher in AD samples compared to control (Fig. 4E). Interestingly, a more stringent analysis of the top 5 percentiles of EVs revealed that the majority of AT8 detections are found in EVs in the 50-200 nm diameter range, which encompasses >90% of the EVs in our samples (Fig. 1 and Fig. 4B). When the mean detections per EV were plotted in this EV size range, a sharp peak at 100 nm was observed for AD samples, suggesting that this population is specific to pathological Tau-carrying EVs (Fig. 4F).

Taken together, our data show that SMLM enables a highly sensitive analysis of the EV cargo of single vesicles and at population level. The results demonstrate that AD brain-derived EVs have a significantly higher AT8-positive Tau cargo than control samples, both in qualitative and in quantitative terms, as judged by a higher ratio of AT8-positive EVs as well as a strongly increased number of AT8 detections per vesicle.

## Discussion

Making use of the outstanding sensitivity and high spatial resolution of single molecule localisation microscopy (SMLM) we have identified pathological Tau proteins in extracellular vesicles (EVs) from human patients with Alzheimer disease. The ability to capture and visualize thousands of vesicles at once ensured statistically meaningful/relevant analysis of EV samples. Our data *i)* demonstrate that AT8-positive EVs represent a relatively large fractions of the entire EV population in AD brains compared to control samples, *ii)* confirm the intraluminal topology of AT8-positive Tau aggregates, and *iii)* provide proof that SMLM analysis of brain-derived EVs can discriminate between AD patients and control individuals.

Key to our study was the ability to isolate intact EVs from post-mortem human brain tissue, which is particularly challenging considering that the varying delays before the tissue is processed and the duration of storage at -80°C may affect the integrity and properties of the EVs(Crescitelli et al. 2025). First, the tissue had to be enzymatically and/or mechanically dissociated to release EVs from the extracellular matrix, while minimizing contamination by intracellular components, including intraluminal vesicles, and alterations of the EV surface epitopes. Following earlier protocols(Crescitelli et al. 2021; Matamoros-Angles et al. 2024; Oosterlynck et al. 2025), we used collagenase D for dissociation of brain tissue due to its very low proteolytic activity. To prevent potential tissue damage, further precautions were taken, such as a short incubation time with the enzyme, limited slicing, and reduced mechanical processing. Secondly, EVs had to be isolated from the resulting interstitial fluid. This can be achieved through a variety of methods, such as differential and density gradient ultracentrifugation, size-exclusion chromatography (SEC), precipitation, and combinations thereof, each of which have their own advantages and limitations. For instance, ultracentrifugation and precipitation-based methods can alter the integrity of the EVs and drastically increase the occurrence of multilamellar and aggregated vesicles as well as contamination by non-vesicular particles(Welsh et al. 2024; Crescitelli et al. 2025) (and our unpublished data). The choice of SEC as a single-step method of EV isolation ensured their good preservation while effectively eliminating most contaminants. This is reflected in the well preserved morphology of the EVs seen in electron micrographs (Fig. 1, Supplementary Fig. S1) as well as in the reconstructed single molecule super-resolution images (Fig. 2, Fig. 3, Fig. 4 and Supplementary Fig. S5).

Up until now, the association of Tau with EVs was demonstrated mainly biochemically by Western blot, enzyme-linked immunosorbent assay (Elisa) or mass spectrometry-based proteomics(Crotti et al. 2019; Fowler et al. 2025; Leroux et al. 2022; Miyoshi et al. 2021; Ruan et al. 2021; Saman et al. 2012; Tyagi et al. 2026). The presence of Tau within the lumen rather than at the surface of the EVs was essentially demonstrated by protease protection assays(Leroux et al. 2022; Ruan et al. 2021). All these experiments were done on bulk EV populations or grossly separated EV subpopulations (e.g. small versus large EVs). Immunogold labeling and transmission EM confirmed the presence of Tau in EVs with single vesicle resolution(Dujardin et al. 2014) (Supplementary Fig. S2), and both patient and recombinant Tau filaments were observed in EVs by cryo TEM(Fowler et al. 2025; Karabova et al. 2026). These techniques are time-consuming, laborious and not quantitative, making them incompatible with high throughput analysis. We therefore implemented dual-colour SMLM as a more direct approach to identify the presence of pathological Tau in patient EVs. The main caveats of SMLM recordings are the repetitive blinking of fluorophores that hampers quantitative analysis and the presence of autofluorescence background that can produce detection artifacts(Camuso et al. 2026). These can be partially eliminated during analysis. By imposing a stringent 25 nm threshold on the localization precision as well as tight intensity ratios for spectral demixing, the number of inaccurate detections becomes negligible (Fig. 2C, Supplementary Fig. S3). We did not correct for multiple detections of the same fluorophores in sequential image frames, because we prioritized the positive identification of Tau aggregates over their exact quantification. Limiting the analysis of Tau signals to clusters of tetraspanin detections, that is to say EVs, provided an additional, spatial filter of the SMLM data. Even so, the strongest argument for the presence of pathological Tau stems from the parallel imaging of thousands of EVs yielding statistically significant datasets (Fig. 3F).

The high spatial resolution of SMLM makes it possible to visualize the ultrastructure of individual EVs as well as their cargo(Alexandre et al. 2025). EVs generally appear as round clusters of detections of varying sizes in super-resolution image reconstructions (Fig. 2, Supplementary Fig. S3). Their size distribution is skewed towards small EVs, with a median value of approximately 80±5 nm. (Fig. 4A,B). This could be a slight overestimate due to the linkage errors caused by antibody labelling (see below) and because the clustering with DBSCAN is inclusive rather than based on the full width at half maximum (FWHM). Nonetheless, the size distribution of EVs recorded by SMLM closely matches the one obtained by EM (Fig. 1, Fig. 4A,B). Both small and large EVs were found to be positive for tau (Fig 4D), in agreement with previous observations(Dujardin et al. 2014; Fowler et al. 2025; Leroux et al. 2022; Oosterlynck et al. 2025; Ruan et al. 2021). Importantly, a peak of the number of Tau detections per vesicle was observed in the 90-130 nm-sized EVs from AD samples but not in control EVs, suggesting that this population is specifically involved in seed propagation (Fig. 4F). In many instances, AT8 detections formed globular or, less frequently, elongated clusters (Fig. 4C, see also Supplementary Fig. S5). These are reminiscent of the small oligomeric or fibrillar forms of Tau, respectively, that have previously been shown to associate with EVs(Fowler et al. 2025; Ruan et al. 2021). To better discriminate different Tau species present in EVs by SMLM will require methodological improvements, such as using smaller fluorophore-conjugated nanobodies against Tau to reduce the linkage errors, or three-dimensional super-resolution imaging to reconstruct the spatial organization of Tau aggregates within the spherical shape of the EVs(Alexandre et al. 2025). Even though the membrane topology of EV-associated tau aggregates cannot be unambiguously determined from our dual-colour SMLM images, their preferential detection in permeabilized EVs argues for a localization of the Tau cargo inside the lumen of EVs from AD brains (Fig. 3C,F).

To what extent the approach presented here can offer new diagnostic solutions for tauopathies depends on the choice of patient samples and the sensitivity of the approach. By analyzing cortical brain tissue of patients in the late stages of AD (Braak V/VI), at which point the pathology is likely to have spread extensively throughout the neocortex(Braak et Braak 1991), we assumed that the proportion of EVs carrying Tau seeds would facilitate detection by SMLM. Indeed, as many as 10% of EVs from AD brains may be positive for AT8 according to our estimates (Fig. 3F). The proportion of Tau-positive EVs is likely to be lower in other brain regions or at earlier disease stages. Future developments need to address the question whether EVs in biological fluids such as plasma or cerebrospinal fluid could be used as diagnostic and prognostic markers. The capacity of SMLM to identify rare, theoretically single target molecules will need to be matched by similar improvements in labelling specificity to avoid false positive detections.

It is tempting to speculate that the selective capture and SMLM analysis of EV subpopulations using specific surface markers will allow us to categorize and quantify the association of Tau seeds more globally. This approach could be extended to other neurodegenerative diseases, since EVs are thought to be involved in the prion-like transmission of other pathological proteins, such as α-synuclein in Parkinson disease or amyloid β in AD(Chen et al. 2025; Colin et al. 2020).

## Materials and Methods

### Samples

Human post-mortem brain tissue samples from Alzheimer’s disease and control subjects (Supplementary Table S1) were obtained through a brain donation program of the brain bank Neuro-CEB (GIE Neuro-CEB BB-0033-00011) at the Hôpital de la Pitié-Salpétrière (Paris, France). The consent forms were signed by the patients themselves, or their next of kin in their name, in accordance with French bioethics laws. All the experiments were done according to the legislation in force for the use of biological samples in scientific research and respecting the anonymity of the donors.

### Brain-derived EV isolation

Frozen brain tissues (0.5-1 g) were gently cut on ice into ∼5 mm^3^ pieces, and then dissociated using collagenase D (2 mg/mL) and DNase I (40 U/mL) in Hibernate-E medium containing 10 mM HEPES-OH pH 7.4, at a ratio of 0.25 g of tissue/mL. After 10 minutes at 37°C, the reaction was stopped on ice by adding 250µL/mL of a 5X quenching buffer (1 tablet each of Complete mini, EDTA-free, protease inhibitor cocktail and PhosSTOP phosphatase inhibitor cocktail, 5 mM PMSF, 2 mL of HEPES-buffered saline (10 mM HEPES pH 7.4, 137 mM NaCl, 5 mM KCl)). The tissues were further dissociated by gently pipetting up and down up to ten time. Then, tissue pieces and debris were eliminated by low-speed centrifugations, once at 500 *g* for 5 min, and twice at 2000 *g* for 10 min. The 2000 *g* supernatant was then filtered using 0.45 μm syringe-filters and concentrated to a final volume of ∼200 µL by ultrafiltration using 100 kDa MWCO Amicon Ultra centrifugal filters that were pre-washed with HEPES-buffered saline. To preserve the integrity of the EVs, the speed of the centrifugation was kept below 2,500 *g* during the concentration step. To purify EVs, 150 µL of concentrated samples were loaded on Izon qEVsingle (gen 2) size exclusion chromatography columns with a pore size of 35 nm that were equilibrated in PBS at room temperature. Fractions (170 µL each) were collected using an Izon automatic fraction collector (V2), and fractions 2 and 3, which contain the EVs, were pooled, aliquoted, flash-frozen and kept at -80°C until use.

### Electron microscopy

Electron microscopy analysis of EV preparations were done essentially as described previously(Kabani et al. 2020; Kabani et Melki 2015). Briefly, EV preparations were thawed, mixed with an equal volume of 4% paraformaldehyde (PFA) solution in PBS, and incubated for 20 min at 4°C. Fixed EV were then applied to formvar/carbon-coated EM copper grids (Electron Microscopy Sciences) and allowed to adsorb for 20 min at room temperature. EM grids were then washed by sequential transfers on drops of PBS, followed by sequential transfers on drops of distilled water. For immunogold labelling, fixed EVs adsorbed on EM grids were permeabilized in blocking buffer (PBS, 5% BSA) containing 0.02% Triton X-100 for 10 min at room temperature. EM grids were transferred to a drop of blocking buffer for 20 min at room temperature, and then incubated for one hour in PBS containing 1% BSA and recombinant rabbit scFv-Fc AT8 antibodies against Tau (a gift from Dr Luc Bousset; at a 1:100 dilution). EM grids were then washed in multiple drops of PBS containing 0.5% BSA and then incubated for 30 min at room temperature in the same buffer containing gold-conjugated secondary antibodies (at a 1:50 dilution; Aurion). EM grids were then washed in drops of PBS, fixed with 1% glutaraldehyde for 5 min at room temperature, and then further washed in multiple drops of distilled water. Following negative-staining with 1% uranyl acetate for 10 min at room temperature, samples were imaged in a Jeol 1400 transmission electron microscope (Jeol, Croissy/Seine, France). Images were recorded with a Gatan Orius CCD camera (Gatan, Pleasanton, CA). The size distributions of EVs was determined from these electron micrographs using the Image J software (NIH) and by two operators, each of whom measured the diameter of at least 200 EVs from 5 different field of views. Data were compiled and analyzed using Graphpad Prism software (version 11.0.0).

### Western blots and dot blots

Western blotting analyses were performed using standard procedures(Kabani et al. 2014, 2011). Equal volumes of EVs were lysed with 1X RIPA buffer (SERVA Electrophoresis) at a 5:1 ratio for 10 min on ice, and then dissolved in 4X Laemmli sample buffer (BioRad) under non-reducing (for CD63 and CD81) or under reducing (for CD9 and Hsc70) conditions and separated by SDS-PAGE. Following electrophoresis, samples were transferred on nitrocellulose membranes. For dot blots, the same amounts of EVs (6.4×10^7^ particles in a total volume of 4 µL) were spotted on nitrocellulose membranes until drying. Membranes were then saturated with 5% skimmed milk in Tris-Buffered Saline containing 0.1% Tween-20 (TBST). Target proteins were visualized after incubation with primary antibodies (α-CD63 (1:1000; Abcam Ab315108), α-CD81 (1:1000; SantaCruz sc-23963 5A6), α-CD9 (1:1000; Abcam ab307085)), α-Hsc70 (1:1000; SantaCruz sc-7298 B-6), recombinant rabbit scFv-Fc AT8 antibodies against Tau (1:500) followed by secondary antibodies coupled to HRP (Invitrogen; 1:5000). Membranes were developed using Super Signal West Femto enhanced chemiluminescence reagents (Thermo Fisher Scientific), imaged with an ImageQuant LAS 4000 (GE Healtcare) imaging system and, when indicated, quantified with ImageJ (NIH).

### Tunable Resistive Pulse Sensing (TRPS)

The concentration and size distribution of EV preparations were measured using an Exoid TRPS instrument (IZON Science) according to the manufacturer’s instructions. We used NP200 nanopores that cover an analysis range of particles from 85 to 500 nm. Calibration particles (CPC200) and EV samples were diluted in freshly filtered (0.22 µm) PBS. Two different dilutions were measured for each sample. All measurements were performed by applying a pressure of 500 Pa and adjusting the nanopore stretch (typically 47-49 mm) and voltage to achieve a current of 120-140 nA. Data acquisition and analysis were performed using the Izon Data Suite software (v1.0.2.32).

### Single Molecule Localization Microscopy

#### Sample preparation for SMLM

Purified EVs were immobilized on glass coverslips and labelled using the Smart EVs kit (Abbelight) according to the manufacturer’s instructions and with additional changes for double detections. Briefly, 6-well microfluidic chambers (Ibidi) were attached to biotinylated glass coverslips (type 1.5). Each channel was functionalized with biotinylated lectins using a streptavidin linker, and then incubated with blocking buffer (Smart EVs kit; Abbelight). EV samples (at a concentration of 5×10^9^ particles/mL) were applied to each channel and incubated for 2 h at room temperature, then overnight at 4°C, before each channel was rinsed three times with washing buffer. When indicated, the captured EVs were fixed with a 2% paraformaldehyde solution in PBS for 10 min at room temperature, quenched with 50 mM glycine in PBS for 5 min, and permeabilized using 0.02% Triton X-100 in PBS for 10 min at room temperature. EVs were labelled with a mix of AF647-conjugated mouse monoclonal antibodies against human CD9, CD63 and CD81 (Smart EVs kit, Abbelight) for 2 h at room temperature and in the dark. Alternatively, rabbit monoclonal antibodies against individual tetraspanins were used (anti-CD9, ab307085; anti-CD63, ab315108; anti-CD81, ab109201, Abcam; 1:500). For double labelling of Tau protein, samples were further incubated with recombinant rabbit scFv-Fc AT8 antibody (a gift from Luc Bousset, CNRS) for 2 h at room temperature. CF680-coupled secondary antibodies (AB_2535737, Thermo Fisher Scientific, 1:1000) were then applied for 2 h at room temperature and in the dark. Prior to SMLM imaging, the samples were incubated in dSTORM buffer (Abbelight).

#### SMLM acquisition

Labelled IBIDIs slides were placed on the motorized stage of an inverted microscope (Zeiss Axio Observer.Z1 SR) and imaged with a 100x oil immersion objective (Zeiss Plan-Apochromat NA 1.46). SMLM was done with a SAFe 360 nanoscope (Abbelight) installed on the Zeiss setup as described previously (Camuso et al. 2026). For spectral demixing, we employed a 700 nm dichroic mirror to separate the emission of far red fluorophores (AF647 and CF680) into a transmitted channel (>700 nm) and a reflected channel (<700 nm), which were detected with two identical sCMOS cameras (Hamamatsu Orca-Fusion BT, CamT and CamR) with a resulting image pixel size of 97 nm. For each sample, three movies of 20000 frames of 512 x 512 pixels (∼2500 µm^2^) were acquired in streaming mode with an exposure time of 50 ms. A 640 nm excitation laser (Oxxius LPX-640-500) was used at a maximal irradiance of 6.4 kW/cm^2^. To maintain a more or less constant sampling rate, 405 nm laser illumination (irradiance ≤ 23 W/cm^2^) was added incrementally during the recording.

#### Image processing and data analysis

The raw SMLM movies were first processed with Neo analysis software (Abbelight). Single molecule detection, 2D Gaussian fitting, x/y drift correction, and spot alignment were carried out with default settings, resulting in 2D coordinate tables of fluorophore signals that were simultaneous detected with both the transmitted and the reflected cameras (CamT, CamR). The uncertainty threshold for the localization precision was set to 25 nm. The intensity ratio of each fluorophore detection in the two channels (I_T_/(I_T_ + I_R_)) was calculated, and spectral demixing applied using the range of 0.18–0.42 for AF647 and 0.48–0.74 for CF680. The demixed detections in the AF647 channel were clustered using DBSCAN by setting the radius ε=80 nm and the minimum number of neighbors to n=10. For quantitative analysis the detections in both channels were rendered and exported as super-resolution tiff images with a pixel size of 20 nm (uniform Gaussian). Single detections in the AF647 image were removed using an intensity filter set at 110 arbitrary units (a.u.) in Fiji. After filling possible holes of the EV clusters, the objects were segmented, and the intensities of each cluster measured in the AF647 channel as well as in the corresponding CF680 image. The resulting intensities were divided by 1032 a.u. to convert them into detection numbers.

### Statistical analysis

All statistical analysis was performed using Graphpad Prism software (version 11.0.0). The statistical tests used are indicated in the figure legends and results section. Data are expressed as mean ± SEM (standard error of the mean) unless indicated otherwise.

## Data availability

All analysed data are included in this article and its supplementary information files. Raw data will be submitted to a data repository upon article acceptance.

## Author contributions

MK and CGS conceived and supervised the study and acquired funding. DG and AB carried out the experiments and analysed the data with input from MK and CGS. All authors participated in the writing of the manuscript.

## Acknowledgements

We thank the donors and the brain bank Neuro-CEB Neuropathology Network supported by patients’ associations and research foundations (ARSLA, France DFT, Fondation Vaincre Alzheimer, France Parkinson, Fondation ARSEP, CSC). We further thank Luc Bousset for the generous gift of antibodies, Serena Camuso and Clémence Mille for technical assistance, and Caterina Severi and Cataldo Schietroma (Abbelight) for sharing their expertise in SMLM analysis of EVs.

## Note

The NeuroCEB Neuropathology network includes: Dr Franck Letournel (CHU Angers), Dr Marie-Laure Martin-Négrier (CHU Bordeaux), Pr Françoise Chapon (CHU Caen), Pr Catherine Godfraind (CHU Clermont-Ferrand), Pr Claude-Alain Maurage (CHU Lille), Dr Vincent Deramecourt (CHU Lille), Dr David Meyronnet (CHU Lyon), Dr Nathalie Streichenberger (CHU Lyon), Dr André Maues de Paula (CHU Marseille), Pr Valérie Rigau (CHU Montpellier), Dr Fanny Vandenbos-Burel (Nice), Pr Charles Duyckaerts (CHU PS Paris), Pr Danielle Seilhean (CHU PS, Paris), Dr Susana Boluda (CHU PS, Paris), Dr Isabelle Plu (CHU PS, Paris), Dr Serge Milin (CHU Poitiers), Dr Dan Christian Chiforeanu (CHU Rennes), Pr Annie Laquerrière (CHU Rouen), Dr Béatrice Lannes (CHU Strasbourg).

## Supplementary data

**Supplementary Figure S1. Electron microscopy analysis of purified brain-derived EVs.** EVs isolated by size exclusion chromatography (SEC) from **(A)** AD or **(B)** control tissues were visualized by negative-staining transmission electron microscopy (TEM). Representative panels of EVs of varying sizes (each from the same individual) are shown (scale bar: 200 nm).

**Supplementary Figure S2. Immunogold electron microscopy analysis of AD brain-derived EVs.** AD brain-derived EVs were fixed with 2% paraformaldehyde, adsorbed onto electron microscopy (EM) grids, and permeabilized with 0.02% Triton X-100 for 10 min. Immunogold labeling of Tau was then performed using AT8 primary antibodies and secondary gold-conjugated antibodies. Electron microscopy grids were fixed with 1% glutaraldehyde for 5 min at room temperature and visualized by the negative-stain EM (scale bars, 100 nm; arrowheads point to gold-labeled AT8-positive Tau within vesicles).

**Supplementary Figure S3. Dual-colour SMLM of brain-derived EVs.** Single molecule imaging was performed on brain-derived EVs labelled with CD9, CD63 and CD81 coupled to either AF647 **(A)** or CF680 **(B)**. **(A,B)** Left: the intensity ratio histogram of single molecule detections is shown in gray, as well as the intensity ranges chosen to separate the AF647 detections (red, 0.18-42) and the CF680 detections (cyan, 0.48-0.74). Note that no signals are detected in the CF680 channel when AF647 is used to label tetraspanins and vice-versa, attesting to the specificity of fluorophore attribution by spectral demixing. Right: Super-resolution pointillist images of representative EVs labeled with either fluorophore (scale: 100 nm).

**Supplementary Figure S4. Electron microscopy analysis of fixed and permeabilized purified brain-derived EVs.** Electron micrographs of brain-derived EVs that were adsorbed on EM grids and that were left untreated (upper panels), or fixed with 2% PFA in PBS for 10 min and permeabilized with 0.02% Triton X100 for 10 min (lower panels, scale bars: 200 nm). No obvious differences in morphology were seen.

**Supplementary Figure S5. Single molecule super-resolution imaging of purified EVs.** Dual-colour SMLM was carried out with EVs labelled with anti-CD9/63/81 (AF647, red) and anti-Tau antibodies (AT8, CF680, cyan), with or without fixation and permeabilization (see Fig. 4). Representative pointillist images of single EVs from one control and one AD brains are shown.

**Supplementary Table S1.** Human tissues used in this study

## References

1. Alexandre, Lucile, Daniele D’Arrigo, Nicolas Kuszla, et al. 2025. « Illuminating Extracellular Vesicles Biology with Super-Resolution Microscopy: Insights into Morphology and Composition ». ACS Nano 19 (27): 24148–73. 10.1021/acsnano.5c00380.

2. Arriagada, Paulina V., John H. Growdon, E. Tessa Hedley-Whyte, et Bradley T. Hyman. 1992. « Neurofibrillary tangles but not senile plaques parallel duration and severity of Alzheimer’s disease ». Neurology 42 (3): 631–39. 10.1212/wnl.42.3.631.

3. Braak, H., et E. Braak. 1991. « Neuropathological stageing of Alzheimer-related changes ». Acta Neuropathologica 82 (4): 239–59. 10.1007/BF00308809.

4. Buzas, Edit I. 2023. « The Roles of Extracellular Vesicles in the Immune System ». Nature Reviews Immunology 23 (4): 236–50. 10.1038/s41577-022-00763-8.

5. Camuso, Serena, Yana Vella, Souad Youjil Abadi, Clémence Mille, Bert Brône, et Christian G. Specht. 2026. « Single Molecule Counting Detects Low-Copy Glycine Receptors in Hippocampal and Striatal Synapses ». eLife 14 (février): RP109447. 10.7554/eLife.109447.

6. Chastagner, Patricia, Frida Loria, Jessica Y. Vargas, et al. 2020. « Fate and Propagation of Endogenously Formed Tau Aggregates in Neuronal Cells ». EMBO Molecular Medicine 12 (12): e12025. 10.15252/emmm.202012025.

7. Chen, Jun, Chen Tian, Xiao Xiong, Ying Yang, et Jing Zhang. 2025. « Extracellular vesicles: new horizons in neurodegeneration ». eBioMedicine 113 (mars). 10.1016/j.ebiom.2025.105605.

8. Colin, Morvane, Simon Dujardin, Susanna Schraen-Maschke, et al. 2020. « From the prion-like propagation hypothesis to therapeutic strategies of anti-tau immunotherapy ». Acta Neuropathologica 139 (1): 3–25. 10.1007/s00401-019-02087-9.

9. Crescitelli, Rossella, Yiyao Huang, An Hendrix, et al. 2025. « Recommendations for Studying In Situ Extracellular Vesicles From Solid Tissue ». Journal of Extracellular Vesicles 14 (11): e70185. 10.1002/jev2.70185.

10. Crescitelli, Rossella, Cecilia Lässer, et Jan Lötvall. 2021. « Isolation and characterization of extracellular vesicle subpopulations from tissues ». Nature Protocols 16 (3): 1548–80. 10.1038/s41596-020-00466-1.

11. Crotti, Andrea, Hameetha Rajamohamend Sait, Kathleen M. McAvoy, et al. 2019. « BIN1 Favors the Spreading of Tau via Extracellular Vesicles ». Scientific Reports 9 (1): 9477. 10.1038/s41598-019-45676-0.

12. Dujardin, Simon, Séverine Bégard, Raphaëlle Caillierez, et al. 2014. « Ectosomes: A new mechanism for non-exosomal secretion of Tau protein ». PLoS ONE 9 (6). 10.1371/journal.pone.0100760.

13. Fitzpatrick, Anthony W. P., Benjamin Falcon, Shaoda He, et al. 2017. « Cryo-EM structures of tau filaments from Alzheimer’s disease ». Nature 547 (7662): 185–90. 10.1038/nature23002.

14. Fowler, Stephanie L., Tiana S. Behr, Emir Turkes, et al. 2025. « Tau filaments are tethered within brain extracellular vesicles in Alzheimer’s disease ». Nature neuroscience 28 (1): 40–48. 10.1038/S41593-024-01801-5.

15. Frost, Bess, Rachel L. Jacks, et Marc I. Diamond. 2009. « Propagation of Tau Misfolding from the Outside to the inside of a Cell ». The Journal of Biological Chemistry 284 (19): 12845–52. 10.1074/jbc.M808759200.

16. Gauthier, Sébastien A., Rocío Pérez-González, Ajay Sharma, et al. 2017. « Enhanced Exosome Secretion in Down Syndrome Brain - a Protective Mechanism to Alleviate Neuronal Endosomal Abnormalities ». Acta Neuropathologica Communications 5 (1): 65. 10.1186/s40478-017-0466-0.

17. Goedert, M., R. Jakes, R. A. Crowther, et al. 1993. « The Abnormal Phosphorylation of Tau Protein at Ser-202 in Alzheimer Disease Recapitulates Phosphorylation during Development ». Proceedings of the National Academy of Sciences of the United States of America 90 (11): 5066–70. 10.1073/pnas.90.11.5066.

18. Goedert, M., R. Jakes, et E. Vanmechelen. 1995. « Monoclonal Antibody AT8 Recognises Tau Protein Phosphorylated at Both Serine 202 and Threonine 205 ». Neuroscience Letters 189 (3): 167–69. 10.1016/0304-3940(95)11484-e.

19. Holmes, Brandon B., Sarah L. DeVos, Najla Kfoury, et al. 2013. « Heparan sulfate proteoglycans mediate internalization and propagation of specific proteopathic seeds ». Proceedings of the National Academy of Sciences of the United States of America 110 (33). 10.1073/pnas.1301440110.

20. Johnstone, R. M., M. Adam, J. R. Hammond, L. Orr, et C. Turbide. 1987. « Vesicle Formation during Reticulocyte Maturation. Association of Plasma Membrane Activities with Released Vesicles (Exosomes) ». The Journal of Biological Chemistry 262 (19): 9412–20.

21. Kabani, Mehdi, Bruno Cosnier, Luc Bousset, Jean Pierre Rousset, Ronald Melki, et Céline Fabret. 2011. « A mutation within the C-terminal domain of Sup35p that affects [PSI+] prion propagation ». Molecular Microbiology 81 (3): 640–58. 10.1111/j.1365-2958.2011.07719.x.

22. Kabani, Mehdi, et Ronald Melki. 2015. « Sup35p in its soluble and prion states is packaged inside extracellular vesicles ». mBio 6 (4). 10.1128/mBio.01017-15.

23. Kabani, Mehdi, Marion Pilard, et Ronald Melki. 2020. « Glucose Availability Dictates the Export of the Soluble and Prion Forms of Sup35p via Periplasmic or Extracellular Vesicles ». Molecular Microbiology 114 (2): 322–32. 10.1111/mmi.14515.

24. Kabani, Mehdi, Virginie Redeker, et Ronald Melki. 2014. « A role for the proteasome in the turnover of sup35p and in [PSI+] prion propagation ». Molecular Microbiology 92 (3): 507–28. 10.1111/mmi.12572.

25. Karabova, Maria Kreger, Anna Del Ser-Badia, Anne Hedegaard, et al. 2026. « Tracking Tau and Cellular Responses in Human iPSC-Microglia: From Uptake to Seedable Secretion, Including in Extracellular Vesicles ». Alzheimer’s & Dementia: The Journal of the Alzheimer’s Association 22 (4): e71337. 10.1002/alz.71337.

26. Kim, Yohan, Rocío Pérez-González, Chelsea Miller, et al. 2022. « Sex Differentially Alters Secretion of Brain Extracellular Vesicles During Aging: A Potential Mechanism for Maintaining Brain Homeostasis ». Neurochemical Research 47 (11): 3428–39. 10.1007/s11064-022-03701-1.

27. Koss, David J., Glynn Jones, Anna Cranston, Heidi Gardner, Nicholas M. Kanaan, et Bettina Platt. 2016. « Soluble Pre-Fibrillar Tau and β-Amyloid Species Emerge in Early Human Alzheimer’s Disease and Track Disease Progression and Cognitive Decline ». Acta Neuropathologica 132 (6): 875–95. 10.1007/s00401-016-1632-3.

28. Krämer-Albers, Eva-Maria. 2022. « Extracellular Vesicles at CNS barriers: Mode of action ». Current Opinion in Neurobiology 75 (août): 102569. 10.1016/j.conb.2022.102569.

29. Leroux, Elodie, Romain Perbet, Raphaëlle Caillierez, et al. 2022. « Extracellular vesicles: Major actors of heterogeneity in tau spreading among human tauopathies ». Molecular Therapy 30 (2): 782–97. 10.1016/j.ymthe.2021.09.020.

30. Malia, Thomas J., Alexey Teplyakov, Robin Ernst, et al. 2016. « Epitope Mapping and Structural Basis for the Recognition of Phosphorylated Tau by the Anti-Tau Antibody AT8 ». Proteins 84 (4): 427–34. 10.1002/prot.24988.

31. Matamoros-Angles, Andreu, Emina Karadjuzovic, Behnam Mohammadi, et al. 2024. « Efficient Enzyme-Free Isolation of Brain-Derived Extracellular Vesicles ». Journal of Extracellular Vesicles 13 (11): e70011. 10.1002/jev2.70011.

32. Matsuo, E. S., R. W. Shin, M. L. Billingsley, et al. 1994. « Biopsy-Derived Adult Human Brain Tau Is Phosphorylated at Many of the Same Sites as Alzheimer’s Disease Paired Helical Filament Tau ». Neuron 13 (4): 989–1002. 10.1016/0896-6273(94)90264-x.

33. Miyoshi, Emily, Tina Bilousova, Mikhail Melnik, et al. 2021. « Exosomal Tau with Seeding Activity Is Released from Alzheimer’s Disease Synapses, and Seeding Potential Is Associated with Amyloid Beta ». Laboratory Investigation 101 (12): 1605–17. 10.1038/s41374-021-00644-z.

34. Niel, Guillaume van, David R. F. Carter, Aled Clayton, Daniel W. Lambert, Graça Raposo, et Pieter Vader. 2022. « Challenges and directions in studying cell-cell communication by extracellular vesicles ». Nature reviews. Molecular cell biology 23 (5): 369–82. 10.1038/S41580-022-00460-3.

35. Oosterlynck, Marie, Elodie Leroux, Balasubramaniam Namasivayam, et al. 2025. « Stratification of Brain-Derived Extracellular Vesicles of Alzheimer’s Disease Patients Indicates a Unique Proteomic Content and a Higher Seeding Capacity of Small Extracellular Vesicles ». Translational Neurodegeneration 14 (1): 63. 10.1186/s40035-025-00519-z.

36. Ruan, Zhi, Dhruba Pathak, Srinidhi Venkatesan Kalavai, et al. 2021. « Alzheimer’s disease brain-derived extracellular vesicles spread tau pathology in interneurons ». Brain 144 (1): 288–309. 10.1093/BRAIN/AWAA376.

37. Saman, Sudad, WonHee Kim, Mario Raya, et al. 2012. « Exosome-Associated Tau Is Secreted in Tauopathy Models and Is Selectively Phosphorylated in Cerebrospinal Fluid in Early Alzheimer Disease * ». Journal of Biological Chemistry 287 (6): 3842–49. 10.1074/jbc.M111.277061.

38. Tyagi, Mitali, Eric de Hoog, Matthew Grega, et al. 2026. « Arc Mediates Intercellular Tau Transmission via Extracellular Vesicles ». Cell, juin 29, S0092-8674(26)00695-1. 10.1016/j.cell.2026.06.008.

39. Vrij, Jeroen de, Sybren L. N. Maas, Malisa van Nispen, et al. 2013. « Quantification of Nanosized Extracellular Membrane Vesicles with Scanning Ion Occlusion Sensing ». Nanomedicine (London, England) 8 (9): 1443–58. 10.2217/nnm.12.173.

40. Wang, Yipeng, Varun Balaji, Senthilvelrajan Kaniyappan, et al. 2017. « The Release and Trans-Synaptic Transmission of Tau via Exosomes ». Molecular Neurodegeneration 12 (1): 5. 10.1186/s13024-016-0143-y.

41. Welsh, Joshua A., Deborah C. I. Goberdhan, Lorraine O’Driscoll, et al. 2024. « Minimal information for studies of extracellular vesicles (MISEV2023): From basic to advanced approaches ». Journal of extracellular vesicles 13 (2): e12404. 10.1002/jev2.12404.

42. Yang, Yang, Diana Arseni, Wenjuan Zhang, et al. 2022. « Cryo-EM structures of amyloid-β 42 filaments from human brains ». Science 375 (6577): 167–72. 10.1126/science.abm7285.

