## Supplementary figures and table for "Single molecule detection of Tau seeds in extracellular vesicles from Alzheimer’s disease brains"

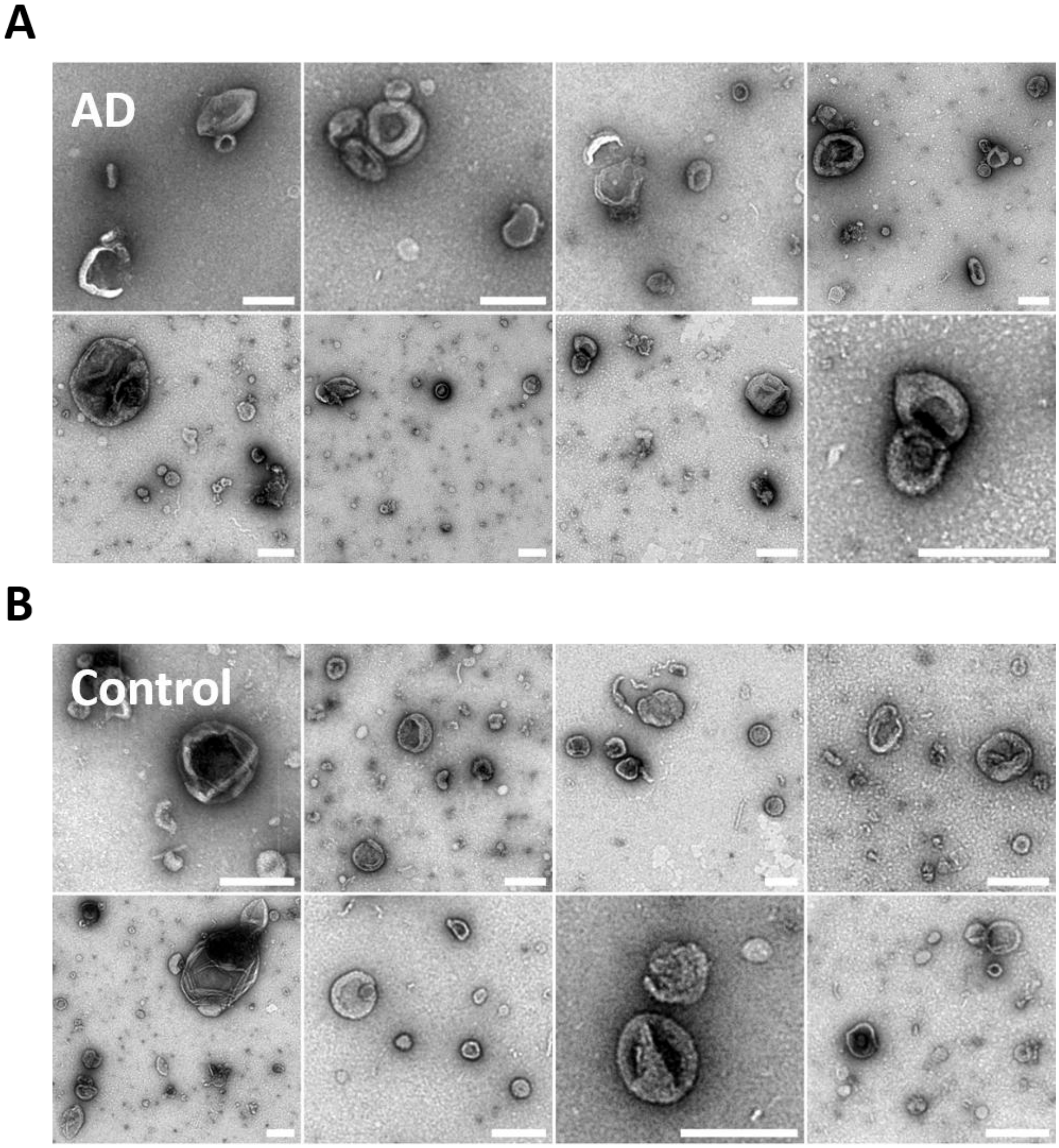


**Supplementary Figure S1. Electron microscopy analysis of purified brain-derived EVs.** EVs isolated by size exclusion chromatography (SEC) from **(A)** AD or **(B)** control tissues were visualized by negative-staining transmission electron microscopy (TEM). Representative panels of EVs of varying sizes (each from the same individual) are shown (scale bar: 200 nm).

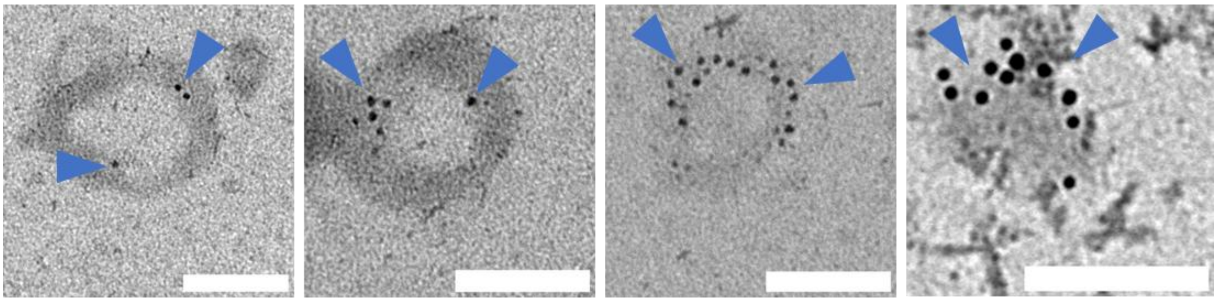

**Supplementary Figure S2. Immunogold electron microscopy analysis of AD brain-derived EVs.** AD brain-derived EVs were fixed with 2% paraformaldehyde, adsorbed onto electron microscopy (EM) grids, and permeabilized with 0.02% Triton X-100 for 10 min. Immunogold labeling of Tau was then performed using AT8 primary antibodies and secondary gold-conjugated antibodies. Electron microscopy grids were fixed with 1% glutaraldehyde for 5 min at room temperature and visualized by the negative-stain EM (scale bars, 100 nm; arrowheads point to gold-labeled AT8-positive Tau within vesicles).

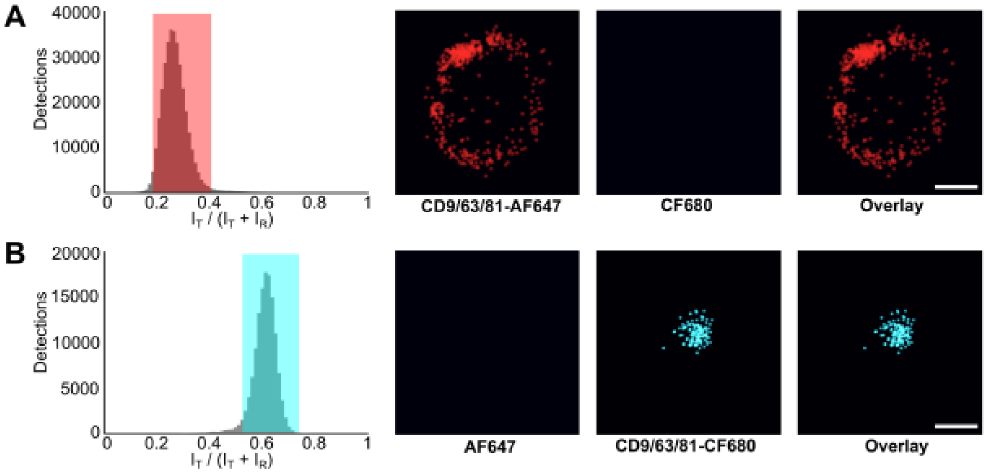


**Supplementary Figure S3. Dual-colour SMLM of brain-derived EVs.** Single molecule imaging was performed on brain-derived EVs labelled with CD9, CD63 and CD81 coupled to either AF647 **(A)** or CF680 **(B)**. **(A,B)** Left: the intensity ratio histogram of single molecule detections is shown in gray, as well as the intensity ranges chosen to separate the AF647 detections (red, 0.18-42) and the CF680 detections (cyan, 0.48-0.74). Note that no signals are detected in the CF680 channel when AF647 is used to label tetraspanins and vice-versa, attesting to the specificity of fluorophore attribution by spectral demixing. Right: Super-resolution pointillist images of representative EVs labeled with either fluorophore (scale: 100 nm).

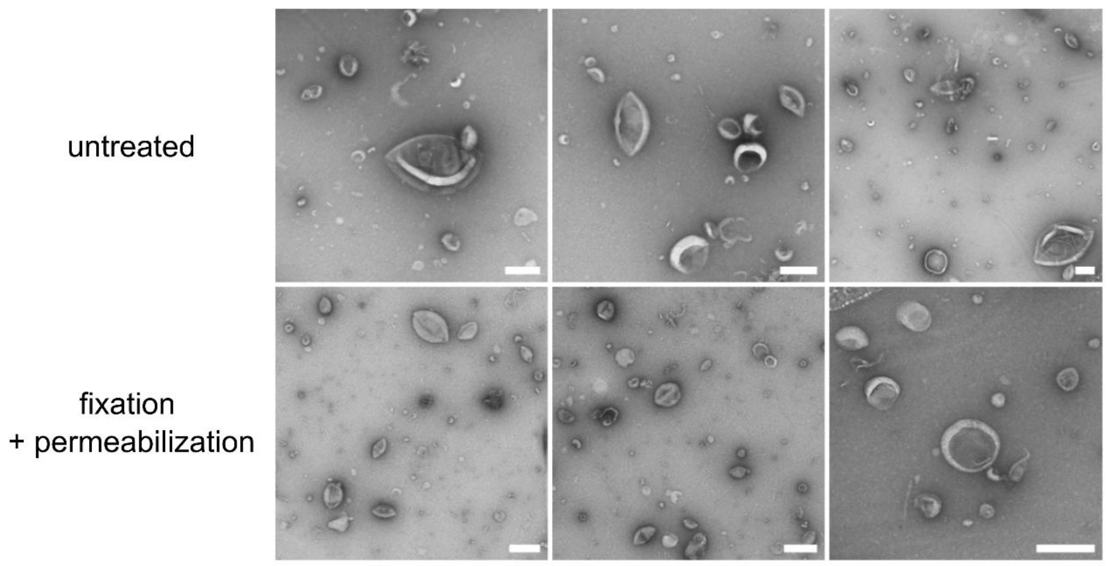


**Supplementary Figure S4. Electron microscopy analysis of fixed and permeabilized purified brain-derived EVs.** Electron micrographs of brain-derived EVs that were adsorbed on EM grids and that were left untreated (upper panels), or fixed with 2% PFA in PBS for 10 min and permeabilized with 0.02% Triton X100 for 10 min (lower panels, scale bars: 200 nm). No obvious differences in morphology were seen.

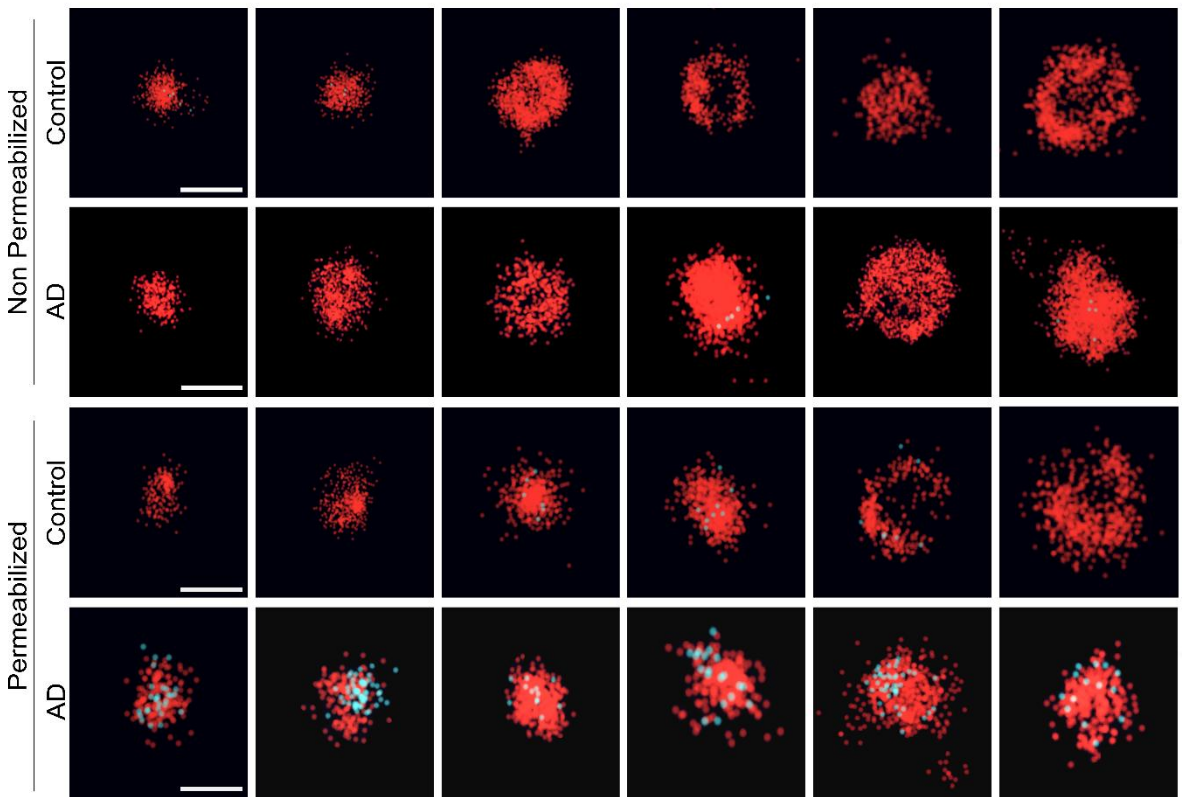


**Supplementary Figure S5. Single molecule super-resolution imaging of purified EVs.**

Dual-colour SMLM was carried out with EVs labelled with anti-CD9/63/81 (AF647, red) and anti-Tau antibodies (AT8, CF680, cyan), with or without fixation and permeabilization (see Fig. 4). Representative pointillist images of single EVs from one control and one AD brains are shown.

**Supplementary Table S1**. Human tissues used in this study

| Patient ID | Sex | Age (years) | Diagnosis | Braak stage | Thal stage |
|---|---|---|---|---|---|
| Alz1 | M | 69 | Alzheimer | VI | 5 |
| Alz2 | M | 78 | Alzheimer | VI | 5 |
| Alz3 | M | 90 | Alzheimer | VI | 4 |
| Alz4 | F | 72 | Alzheimer | VI | 5 |
| Alz5 | F | 72 | Alzheimer | V | 5 |
| Alz6 | F | 64 | Alzheimer | V-VI | N/A |
| Alz7 | M | 82 | Alzheimer | VI | 3 |
| Ctrl1 | F | 76 | Control | 0 | 0 |
| Ctrl2 | F | 92 | Control | III | 0 |
| Ctrl3 | F | 89 | Control | III | 0 |
| Ctrl4 | F | 60 | Control | 0 | 0 |
| Ctrl5 | F | 80 | Control | 0 | 0 |
| Ctrl6 | M | 84 | Control | 0 | 1 |
